# Evidence of kinesin motors involved in stable kinetochore assembly during early meiosis

**DOI:** 10.1101/2022.12.27.522002

**Authors:** Seema Shah, Priyanka Mittal, Deepanshu Kumar, Anjani Mittal, Santanu K Ghosh

## Abstract

The characteristic ‘bi-lobed’ organization of the kinetochores observed during mitotic metaphase is a result of separation of the sister kinetochores into two clusters upon their stable end-on attachment to the microtubules emanating from opposite spindle poles. In contrast, during metaphase I of meiosis despite bi-orientation of the homologs, we observe that the kinetochores are linearly dispersed between the two spindle poles indicating that pole-distal and pole-proximal kinetochores are attached laterally and end-on, respectively to the microtubules. Colocalization studies of kinetochores and kinesin motors suggest that budding yeast kinesin 5, Cin8 and Kip1 perhaps localize to the end-on attached kinetochores while kinesin 8, Kip3 resides at all the kinetochores. Unlike mitosis in budding yeast, in meiosis, the outer kinetochores assemble much later after prophase I. From the findings including co-appearance of kinesin 5 and the outer kinetochore protein Ndc80 at the centromeres after prophase I and a reduction in Ndc80 level in Cin8 null mutant, we propose that kinesin motors are required for reassembly and stability of the kinetochores during early meiosis. Thus, this work reports yet another meiosis specific function of kinesin motor.

## Introduction

The kinetochore, a multi-protein complex formed on the centromeric DNA, is the key player that promotes chromosome segregation by attaching the chromosomes to the microtubules during mitotic and meiotic cell divisions. In meiosis, the kinetochore complex undergoes various organizational rearrangements to promote the meiosis I specific events. For instance, a side-by-side orientation of sister kinetochores in meiosis I, as opposed to back-to-back in mitosis or meiosis II, facilitates the mono-orientation of sister chromatids which is aided by the deposition of meiosis-specific proteins at the kinetochores, viz – Spo13, Mam1 in budding yeast (Miller et al., 2012; Petronczki et al., 2006; Rabitsch et al., 2003; Toth et al., 2000), Moa 1 in fission yeast (Yokobayashi and Watanabe, 2005), and Meikin in metazoans (Kim et al., 2015). Another meiosis-specific kinetochore reorganization is the late formation of a matured kinetochore competent to bind to microtubules during prophase I. In mammals, chickens, and nematodes, outer kinetochore proteins dissociate from the centromeres after completion of mitotic chromosome segregation, and they then reassemble during the next mitotic entry at the time of the establishment of kinetochore-microtubule (KT-MT) attachment (Cheeseman et al., 2008; Hemmerich et al., 2008; Meyer et al., 2015a). However, in budding yeast, kinetochores in mitosis remain assembled and attached to the microtubules throughout the mitotic cell cycle except for a brief period when the centromeres replicate (Kitamura et al., 2007a). In contrast, during meiosis in budding and fission yeasts and in mammals, the complete kinetochore assembly takes place much later after prophase I (Hayashi et al., 1998; Jin et al., 1998; Meyer et al., 2013; Miller et al., 2012; Asakawa et al., 2005; Hayashi et al., 2006; Parra et al., 2009). This delayed kinetochore assembly is unique and essential in meiosis as it is believed, at least in budding yeast, to prevent an early KT-MT attachment which is detrimental to the proper incorporation of the monopolin complex (Kim et al., 2013; Meyer et al., 2015a) required for mono-orientation of the sister chromatids. In budding yeast, two separate mechanisms contribute to the delayed assembly of the kinetochore in meiosis. First, aurora B kinase (Ipl1, budding yeast homolog) mediated shedding of outer kinetochore proteins as the cells enter into meiosis (Kim et al., 2013; Meyer et al., 2013; Meyer et al., 2015). Second, programmed transcription of a long inhibitory isoform of *NDC80* (coding an essential outer kinetochore protein) specifically during meiosis instead of transcription of the functional *NDC80* ORF (Chen et al., 2017; Chia et al., 2017). The expression of the inhibitory form is driven by the formation of the meiosis-specific Ime1-Ume6 transcription factor complex which also shuts down the expression of *NDC80* ORF till prophase I. Whether these two mechanisms act independently or are regulated by a common cue is unknown. Nevertheless, following prophase I, when functional Ndc80 is expressed, it reassembles the outer kinetochore establishing the KT-MT attachment. Owing to the temporal difference in dynamics of kinetochore assembly along with the difference in its composition and spindle attachment pattern between mitosis and meiosis I, the essentiality of several of the proteins targetted at the kinetochore differs between these two cell cycles. For example, in contrast to mitosis, inner kinetochore proteins CENP-C and H3 variant CENP-A in *C. elegans* are dispensable for chromosome segregation and recruitment of the outer kinetochore proteins to the centromeres in meiosis, where the assembly of the outer kinetochore proteins depends on the positions of the cross-overs (Monen et al., 2005). Similarly, in budding yeast, the absence of several proteins of the central kinetochore complex Ctf19 results in high chromosome mis-segregation in meiosis than in mitosis (Mehta et al., 2014, Agarwal et al., 2015). In *S. pombe*, kinesin-8 motors Klp5 and Klp6 localize to the kinetochore (Garcia et al., 2002) and are essential for meiotic but not for mitotic chromosome segregation (West et al.,2001, Erent et al., 2012).

Kinesin motor proteins play a pivotal role in chromosome segregation by contributing to chromosome congression, their alignment to the cell-equatorial plate, spindle formation, and chromosome movements. These functions are disbursed through localization of the motors to the kinetochores, chromosomal arms, spindle, and the spindle poles. In budding yeast there are four non-essential nuclear kinesin motors - two kinesin-5 (Cin8 and Kip1), kinesin-8 (Kip3), and kinesin-14 (Kar3). The functions of these motors belonging to kinesin-5 (Cole et al., 1994; Hagan and Yanagida, 1992; Kapitein et al., 2005; Roof et al., 1992; Sharp et al., 1999a), kinesin-8 (Su et al., 2013) and kinesin-14 (Cai et al., 2009; Fink et al., 2009b; Hepperla et al., 2014; Mountain et al., 1999) families are conserved across the eukaryotes. During chromosome segregation, the kinesin-5 and 8, generally microtubule plus-end directed motors, promote centromere clustering and spindle elongation by cross-linking the parallel and anti-parallel microtubules, respectively (Brust-Mascher et al., 2009; Gardner et al., 2008b; Mayr et al., 2007; Tytell and Sorger, 2006; Wargacki et al., 2010a). Moreover, kinesin-8 with a microtubule depolymerase activity is also involved in the coordinated movement of sister chromatids towards spindle poles during anaphase (Gardner et al., 2008b; Stumpff et al., 2008; Su et al., 2011; Tytell and Sorger, 2006; Wargacki et al., 2010a; West et al., 2002). Notably, kinesin 5 in budding yeast also has depolymerase activity which is believed to promote chromosome congression (Gardner et al., 2008)). Interestingly, recently a minus end-directed motility of kinesin-5 has also been identified that may aid in spindle assembly (Shapira et al., 2017)). On the other hand, kinesin-14, a microtubule minus end-directed motor, is required to capture spindle unattached kinetochores and their lateral sliding along the microtubules towards the spindle poles. (Grissom et al., 2009; Gupta Jr et al., 2006; Mayr et al., 2007, Tanaka et al., 2007). The function of kinesin-14 as microtubule depolymerase is also reported, which helps in a nuclear fusion that demonstrates its role in karyogamy (Meluh and Rose 1990, Sproul et al., 2005).

While the functions of the motors during the mitotic cell cycle are well documented, the literature on meiosis in this regard is scanty. On the other hand, a delayed maturation kinetics of the kinetochores, their unique attachment pattern to the microtubule spindle in meiosis I and two-times spindle assembly/disassembly during meiosis in contrast to mitosis, argues for having novel meiosis-specific functions of the motors. For instance, in budding yeast, the kinesin14 motor Kar3 is essential for meiosis but not mitosis (Carol et al., 1997). Recently, a kinesin 14 motor in fission yeast has been implicated in having specific roles in meiosis I spindle stability (Zheng et al. 2020, Loncar et al. 2020). In this context, earlier, we demonstrated the meiosis-specific roles of kinesin-5 and kinesin 8 in genome stability in budding yeast (Mittal et al., 2020). To further delineate the functions of the kinesin motors in meiosis, particularly during early meiosis when the kinetochores transit from a microtubule unattached to an attached mode, we sought to investigate whether the kinesin 5 and 8 motors promote this transition by influencing kinetochore maturation. Using cell cycle-dependent live cell imaging and biochemical assays, we show that motors, particularly Cin8, is involved in achieving fully assembled kinetochore in meiosis. This study reveals yet another meiosis-specific function of motor evolved to promote meiotic chromosome segregation.

## Materials and Methods

### Yeast strains and media

The strains used in this study were of SK1 background and are listed in Table 1 under ‘supplementary information’. The PCR-based C-terminal protein tagging and gene deletion methods were performed as described elsewhere (Wach., 1996). Transformation of the cells with the PCR cassettes was performed as mentioned previously (Gietz and Schiestl, 2007). During meiosis, cells were synchronized at metaphase I and prophase I by shuffling the endogenous promoters of *CDC20* and *NDT80* with *P_CLB2_* and *P_GAL1_* constructs, respectively, as mentioned earlier (Benjamin *et al*., 2003; Lee and Amon, 2003). For mitotic cell cycle arrest at metaphase, Cdc20 was depleted using auxin-inducible degron system as used before (Nishimura et al., 2009).

For the selection of cells on a dropout media along with the antibiotic G418, instead of ammonium sulfate, monosodium glutamate (Himedia #RM681, India) was used to obtain better sensitivity against G418 as mentioned earlier (Cheng *et al*., 2000).

### Culture conditions

For meiotic synchronization, cells were first patched on YPG (Yeast extract 1%, peptone 2%, glycerol 2%) to restrain the growth of the petite colonies and then transferred to presporulation medium overnight followed by meiotic induction in sporulation medium (0.02% raffinose, 1% potassium acetate) as mentioned earlier (Cha et al., 2000a; Mehta et al., 2014).

For the auxin (3-indole acetic acid, Sigma # I2886, USA) mediated protein degradation, log phase grown cells were treated with 1.5 mM of auxin for 2 h.

For microtubule disruption in prophase I arrested stage (by p*GAL1-NDT80*), cells were harvested after 6 h in sporulation medium (SPM) and resuspended in SPM containing 120 µg/ml benomyl or 0.4% DMSO (Sigma #D8418, USA) for 2 h before they were analyzed for protein localization. For microtubule disruption during metaphase I arrested stage (by *P_CLB2_-CDC20*), cells after 8 h of meiotic induction were treated with benomyl (Sigma #45339, USA) at a concentration of 120 µg/ml or 0.4% DMSO for 1.5 h before they were analyzed for protein localization.

### Fluorescence microscopy

As mentioned earlier, cells were processed for live cell imaging as described before (Mittal et al., 2020). Typically, the cells were fixed with formaldehyde at the final concentration of 5% for 10 min. After washing the cells with 0.1 M phosphate buffer (pH 7.5), images were acquired using AxioObserver Z1 Zeiss (63X 1.4 NA Objective) inverted microscope. The exposure time was kept constant across the samples for different fluorophores for the intensity comparison. Analyses of the images were done using software AxioVs40 V 4.8.2.0. In case of intense CFP signal, the images were acquired using Zeiss LSM 780 - confocal laser scanning microscope equipped with 32 arrays GaAsP detector, and images were analyzed to avoid the bleed-through GFP channel using Zen 2012 software.

### Intensity measurement

For the intensity measurement, Z-stacks with the maximum intensity were merged. For a dispersed signal like Ndc80 the intensity profile was generated, using the Zen software, by drawing a measurement line through the Ndc80 signal and the other line was drawn through the background in the vicinity of the Ndc80 signal. The value obtained for the background signal was averaged and subtracted from the Ndc80 intensity values.. A similar method was followed for generating the intensity profile of Spc42-CFP.

ImageJ software was utilized for fluorescence intensity quantification, as mentioned earlier (Mittal et al., 2020). The integrated intensity of the region of interest of the fluorescent signal was normalized with the background obtained after averaging the integrated values of three random non-fluorescent areas used as backgrounds.

Fluorescent intensity = Fluorescent signal value of the region of interest – background combined value

### ChIP-qPCR and its quantification

Chromatin immunoprecipitation was performed as described previously (Prajapati et al., 2017). Typically, the chromatin was cross-linked with 1% formaldehyde based on the proximity of the protein to the centromere (2h for the outer kinetochore proteins and one h for the inner kinetochore proteins) and resuspended in lysis buffer supplemented with 1X PIC. Initially, the cells were lysed using glass beads and a mini bead beater (BIOSPEC products). Chromatin was fragmented using a Vibra cell sonicator (40% amplitude, 12 cycles - 21 sec ON, 1 min OFF) or Water bath sonicator(Digenode, 30 cycles-30 sec ON, 30 sec OFF). Aliquots were incubated with anti-GFP (5 µg, Roche, # 11814460001, Germany) for GFP or HA11(5ug, Rabbit polyclonal HA11, Abcam, UK) for 6HA epitope overnight, followed by Protein A sepharose beads(GE Healthcare #17-0780-01 USA) at 4°C for 3 h. After sequential washing, samples were incubated at 65°C water bath overnight for de-crosslinking and further treated with Proteinase K(SRL #39450-01-6, India) and glycogen(Roche # 10901393001, Germany) at 42°C for 2 h. Purified samples were used for qPCR analysis using the CFX96 Real-Time system (BioRad) and SYBR green dye (BioRad, USA) for detection. The relative amounts of immunoprecipitated DNA were quantitated using qPCR (Prajapati et al., 2017). Corrections were applied to account for primer efficiencies below 100% (amplification factors < 2.0) using standardization graphs of CT values against dilutions of the input DNA. The amplification factor ε was estimated as 10^(−1/slope) of the regression line, and the primer efficiency E (%) as {[10^(−1/slope)]– 1} x 100. The fraction of immunoprecipitated DNA in a ChIP sample relative to the input DNA was calculated as ε ∧(− Δ CT), Δ CT = CT (ChIP)–[CT (Input)–log (Input dilution factor)]. The primers used for PCR amplification are listed in Table 2 under ‘Supporting Information’.

### Immunoblotting and its quantification

As mentioned earlier, the extraction of whole cell proteins was done by NaOH treatment (Kushnirov, 2000) with some modifications. For meiotic cells, 10 ml of 1 O.D_600_ culture is pelleted down and treated with 0.1 N NaOH for 30 min.

### Indirect immunofluorescence

This was followed as described earlier (Adams and Pringle 1984; Mehta et al., 2014) with some modifications. Typically, the cells were fixed with 4.5% formaldehyde for 2 h at RT. The fixed cells were washed once with PBS and once with spheroplasting buffer (1.2 M sorbitol, 0.1 M phosphate buffer pH 7.5) and were resuspended in the same solution. Cells were spheroplasted using zymolyase 20T (MP Biomedicals # 32092, USA) in the presence of 25 mM β mercaptoethanol for 1 h at 30 °C. The spheroplasts were washed with spheroplasting solution and transferred to poly-L-lysine (Sigma, # P1524, USA) -coated slides having Teflon-coated wells. The spheroplasts were flattened and permeabilized by immersing the slide in methanol for 5 min and acetone for 30 sec. Blocking was done using 10 mg/ml BSA with 5% skim milk in PBS for 15 min. All the primary and secondary antibodies were diluted using antibody dilution buffer (10 mg/ml BSA in PBS). Spheroplasts were incubated with primary antibodies for 1 h followed by washing with PBS several times and were incubated with pre-adsorbed secondary antibodies for 1 h in the dark. After several washes with PBS, samples were incubated with DAPI (1 μ /ml in 0.1 M phosphate buffer, Invitrogen #D1306, USA) for 15 min dark. 90% g glycerol supplemented with 1 mg/ml phenylenediamine(Sigma # 78429, USA) was used as a mounting medium.

Following antibodies were used at indicated dilutions: Rat anti-tubulin antibody (1:5000, Serotec, # MCA78G, UK), Mouse anti-GFP antibody (1:200, Roche, # 11814460001, Germany), TRITC-conjugated goat anti-rat antibody (1:200, Jackson, #115-485-166, USA), DyLight 488-conjugated goat anti-mouse antibody (1:200, Jackson, # 112-025-167, USA).

## Results

### Subcellular localization of the motor proteins during the early phase of meiosis

In mitosis, the recruitment of Cin8 and Kip1 but not Kip3 at the kinetochore is outer kinetochore protein-dependent (Tytell and Sorger, 2006). Since the outer proteins assemble late at the kinetochore in meiosis in contrast to mitosis, it is intriguing to know the temporal order of localization of the motors and their dependency on the outer proteins at the meiotic kinetochore. We analyzed the subcellular localization of Cin8, Kip1, and Kip3 in live meiotic cells that were arrested at prophase I by repressing the early meiotic transcription factor, Ndt80, as described before (Carlile and Amon, 2008). For imaging purposes, the motors (Cin8, Kip1, and Kip3) and the kinetochore marker Ndc10, an inner kinetochore protein, were marked with EGFP, whereas the outer kinetochore (Ndc80) and SPBs (Spc42) were marked with CFP and m-cherry, respectively.

In budding yeast, after 6 h of meiotic induction as the cells enter into prophase I of meiosis, centromeres become declustered presumably due to the shedding of the outer kinetochores and KT-MT detachment (Asakawa *et al*., 2005). Accordingly, we observed a declustered Ndc10 signal (Figure S1) but no Ndc80 signal (Figure 1A) in prophase I arrested cells. However, we observed Cin8 as a clustered dot-like signal (termed focus) in majority (95%) of the cells (Figure 1A). The focus signal of Cin8 localizes next to undivided SPBs (Figure 1A) and at one end of the spindle, as revealed by tubulin immunostaining (Figure 1B). This localization pattern of Cin8 is microtubule-dependent as the Cin8 signal became dispersed after the removal of the microtubules by benomyl (Figure 1C). In the same stage cells, we found Kip1 either along the spindle (54%) or dispersed throughout the nucleus (43%) (Figure 1D) whereas Kip3 was found along the spindle in most of the cells (81%) (Figure 1E). Since cell biological assays were insufficient to infer the centromeric localization of these proteins, we performed chromatin immunoprecipitation (ChIP) assays from the same cells. After comparing their relative enrichment at the centromeres (*CENIII*) we observed that Kip3, but not Cin8 and Kip1, is localized moderately at the kinetochores (Figure 1F) considering a robust centromere localization of the inner kinetochore protein Ndc10. These results led us to conclude that at prophase I, when the outer kinetochore is not assembled at the centromeres, Cin8 and Kip1 cannot be localized at the kinetochores; instead, they remain at SPB and/or along the spindle with a gross nuclear localization. However, Kip3 can be localized at the centromeres and along the spindle in the same cells.

**Figure 1:**
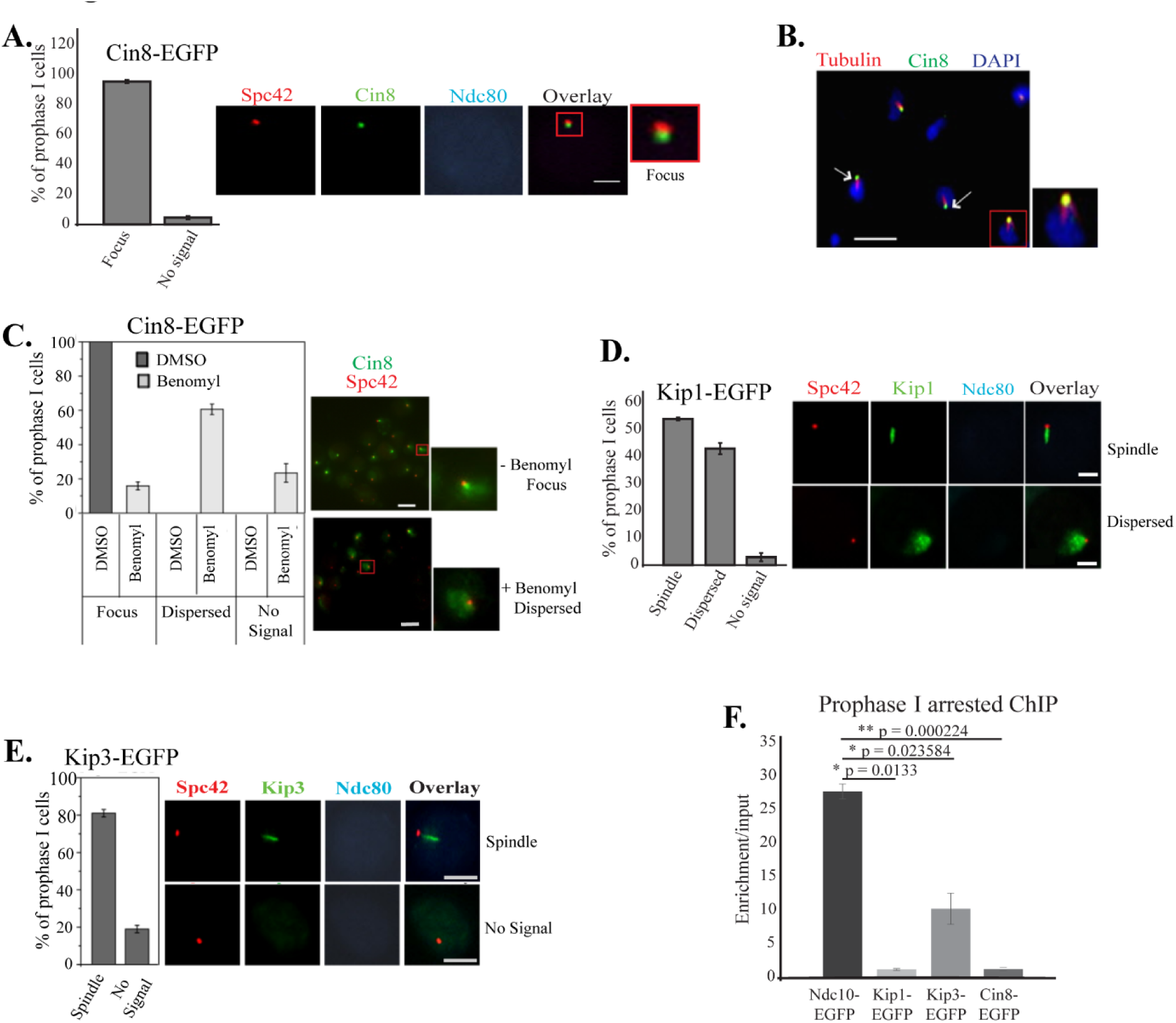
Localization of motor proteins in prophase I cells. Strains harboring Cin8-EGFP (SGY5151) or Kip1-EGFP (SGY5266) or Kip3-EGFP (SGY5272) were arrested at prophase I (by Ndt80 depletion) by releasing the cells into meiosis for 6 h before they were photographed. (A) Right, representative images of live cells showing patterns of Cin8-EGFP localization. Left, the distribution of the patterns amongst the prophase I cells (n = 130) (B) Localization of Cin8-EGFP in cells (SGY5109) containing monopolar spindle. Cells harvested as in A were immunostained using anti-GFP and anti-α-tubulin antibodies. Arrows indicate the localization with respect to the spindle. (C) Right, representative images of live cells showing patterns of Cin8-EGFP localization upon treatment of the prophase I cells with DMSO (mock) or benomyl. Left, the distribution of the patterns in presence (DMSO, n = 117) or absence (benomyl, n = 99) of microtubules. (D and E) Right, representative images of live cells showing patterns of Kip1-EGFP and Kip3-EGFP localization. Left, the distribution of the patterns amongst the prophase I cells (n = 113 for Kip1-EGFP, n = 96 for Kip3-EGFP). ‘n’ represents the total number of cells analysed. (F) ChIP assay using anti-GFP antibodies showing the centromeric localization of the indicated EGFP-fused motor proteins in the prophase I arrested strains Cin8-EGFP (SGY5109), Kip1-EGFP (SGY5220), and Kip3-EGFP (SGY5256). Ndc10-EGFP (SGY5351) was used as a positive control for the assay. The enrichment values of the motor proteins at the *CENIII* locus were plotted. The error bar represents the SEM obtained from two biological replicates. The mentioned p-values were determined with respect to the value obtained for Ndc10. Bar, 2 µm.

### The maintenance of Cin8 and Kip1 at the kinetochore is microtubule-dependent in meiosis

As the cells exit from prophase I and proceed towards metaphase I, the outer kinetochore is recruited at the kinetochore causing its maturation and establishes the connection between the microtubules and kinetochores. Since we wished to investigate the localization behaviour of the motors at the kinetochore during its transition from the immature (unattached) to the mature (attached) state, we next analyzed the metaphase I cells as judged by around 2 µm SPB-SPB distance. Strikingly, unlike mitotic metaphase, we failed to observe the ’bi-lobed’ configuration of the sister kinetochores marked by either Ndc10 or Ndc80; instead, the kinetochores were found linearly dispersed between the two SPBs (Figure. 2A), but not cloud-like dispersed as observed in prophase I (Figure S1). The linearly dispersed pattern indicates that although the kinetochores were mature, all were not yet end-on attached to the microtubules and hence were not fully congressed. We argued that the plausible reason for this could be the non-uniform recruitment of the motors at the kinetochores because in mitosis, it has been shown that Cin8, Kip1, and Kip3 are required at the kinetochores for their clustering to form bi-lobed configuration at metaphase (Gardner et al., 2008b; Tytell and Sorger, 2006; Wargacki et al., 2010a). To verify this possibility, we looked at the motors from the metaphase I cells and found Kip3 is fully colocalized with the linearly dispersed kinetochores (Figure 2B) suggesting that all the kinetochores harbored this motor. However, Cin8 and Kip1 were visualized as a bi-lobed localization between two SPBs and colocalized with a fraction of kinetochores clustered near SPBs (Figure 2B). In mitosis, the kinetochores first attach to the microtubules laterally, which then follow an end-on attachment mediated by outer kinetochore protein Dam1 (Shrestha and Draviam, 2013; Tanaka et al., 2007b). Viewing linearly dispersed kinetochores in metaphase I (Figure. 2A), we speculate that due to late recruitment of the Ndc80 complex at the kinetochore, KT-MT attachment occurs in a stochastic manner in meiosis. In this context, we argue that a fraction of fully mature kinetochores (with both Ndc80 and Dam1 recruited) undergo end-on attachment and form clusters near SPBs. In contrast, the kinetochores that harbour only Ndc80 are laterally attached to the spindle and remain linearly dispersed in between two SPBs. To examine this we marked Dam1 and Ndc80 with EGFP and CFP, respectively in the same cell, where SPB (Spc42) was marked with m-Cherry to judge the cell cycle stages. Cells with SPB-SPB distance around 2 μm suggesting their metaphase stage, were analyzed. As expected, we observed Dam1-EGFP either mostly as one cluster next to each SPB (type I, Figure 3H) or infrequently as linearly dispersed localization in between two SPBs (type II, Figure 3H); whereas in both the types Ndc80-CFP remained linearly dispersed. We presume that the type II category for Dam1-EGFP arises due to asynchronous congression of the end-on attached kinetochores towards the SPBs. As the cell cycle proceeds, all the kinetochores eventually become end-on attached to the microtubules and remain clustered near SPBs during subsequent stages of meiosis (Figure. 2C).

**Figure 2:**
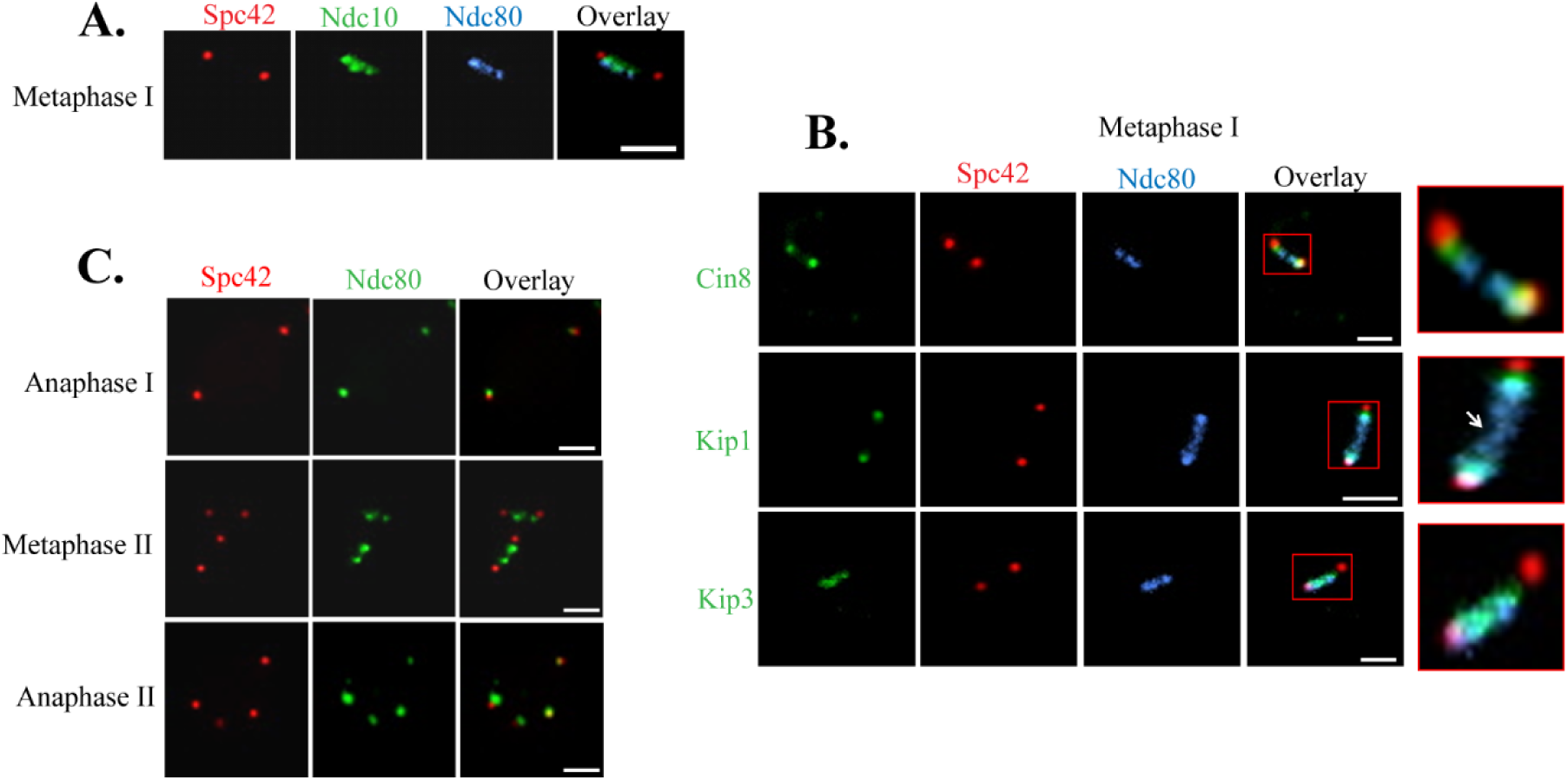
Localization patterns of Cin8 and Kip1 differ from Kip3 in metaphase I cells. (A) Kinetochores are disperesed linearly during metaphase I. Cells (SGY5198) harboring Ndc80-CFP, Ndc10-EGFP, and Spc42-mcherry were released into meiosis for 5-6 h before they were photographed. Representative images show colocalization of Ndc80 and Ndc10 signals. (B) Cin8-EGFP (SGY5070), Kip1-EGFP (SGY5051), and Kip3-EGFP (SGY5165) cells harboring Spc42-mcherry and Ndc80-CFP were similarly treated as in (A). Representative images show a linearly dispersed localization of outer kinetochore protein Ndc80 with colocalized Kip3 while Cin8 and Kip1 are present as clustered foci near SPBs. (C) Representative images show that the Ndc80-EGFP (SGY5130) signal remains clustered during later stages of meiosis after metaphase I. Bar, 2 µm.

**Figure 3:**
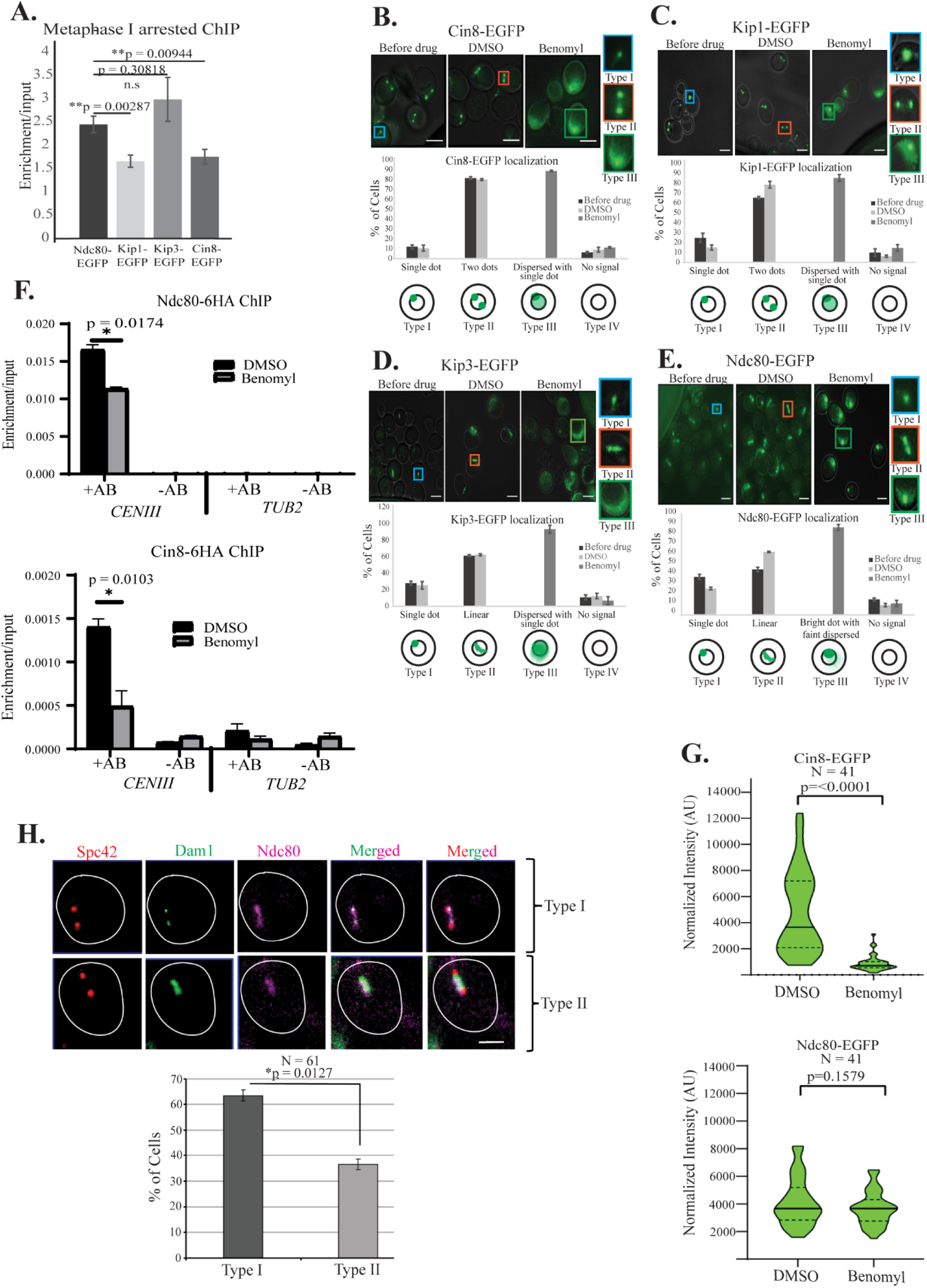
Maintenance of Cin8 and Kip1 at the centromeres during metaphase I is microtubule dependent. (A) ChIP analysis of the strains used in (B-E) shows the association of the indicated proteins with *CENIII*. Ndc80-EGFP was used as a positive control for the centromeric association. The enrichment values of the indicated proteins at the *CENIII* locus were plotted. The error bars represent the SEM obtained from at least two independent replicates. The mentioned p-values were determined between Ndc80 and the respective column bar. (B-E) The localization patterns of Cin8-EGFP (SGY 5119), Kip1-EGFP (SGY 5117), Kip3-EGFP (SGY 5238), and Ndc80-EGFP (SGY 5247) in the presence (DMSO) or absence (benomyl) of microtubules in metaphase I arrested cells. n > 100 for each strain and the error bar represents the mean with SEM from at least two independent experiments. (F) ChIP analysis for localization of Ndc80-6HA (SGY 4206) and Cin8-6HA (SGY 4203) at *CENIII* with (DMSO) or without (Benomyl) microtubules. *TUB2* locus was used as a negative control. The error bars represent the SEM obtained from at least two independent replicates. Bar, 5 µm (G) Quantification of the Cin8-EGFP and Ndc80-EGFP fluorescence intensity for the strain used in B and E, respectively after normalization with the background. (H) Cells (SGY5718) harboring Dam1-EGFP, Ndc80-CFP and Spc42-mcherry were released into meiosis for 6-7 h before they were photographed. Representative images show the localization patterns of the outer kinetochore protein Dam1 either as clustered foci near SPBs (Type I) or as linearly dispersed between two SPBs (Type II) like Ndc80 protein. Bar, 2 µm

Observing the Cin8 and Kip1 signals colocalized with the clustered kinetochores next to the SPBs and Kip3 all along the linearly dispersed kinetochores, we argued that Cin8 and Kip1 require microtubule plus ends engaged at the kinetochores (end-on attachment) to remain associated with the kinetochores, whereas Kip3 may not need any microtubule attachment which is also supported by its localization at the centromeres even in the absence of Ndc80 (Figure. 1F). This implies that Cin8 and Kip1 associate with a subset of the kinetochores attached end-on to the microtubules and remain clustered close to the SPBs, whereas Kip3 and Ndc80 occupy all the kinetochores regardless of the attachment status. In support of this, by ChIP assay, we observed less Cin8 and Kip1 at *CENIII* than Kip3 or Ndc80 (Figure 3A). These observations indicate that perhaps targetting and/or maintenance of Cin8 and Kip1 but not Kip3 and Ndc80 at the kinetochores are microtubule dependent. Since microtubule disruption before or during meiotic S phase arrests cells at G1 or G2, respectively (Hochwagen et al., 2005), we wised to investigate whether the maintenance of Cin8, Kip1, Kip3 and Ndc80 at the kinetochores during metaphase I is microtubule-dependent. We first arrested the cells at metaphase I by Cdc20 depletion (Figure S2A) as described elsewhere (Lee and Amon, 2003) and then removed the microtubules by treating the cells for 1.5 h with benomyl (Figure S2B). We observed that Cin8 and Kip1 showed mostly bi-lobed (two dots) localization in presence of the microtubules (DMSO, type II, Figure 3B,C) as obserbed before (Figure 2B) suggesting that they are at the homolog kinetochore clusters next to the SPBs. Upon removal of the microtubules (Benomyl) they showed mostly dispersed localization along with a single dot within the dispersed stain (type III, Figure 3B,C) suggesting they are mostly dislodged from the kinetochores but some are still at the homolog kinetochores which became coalesced due to removal of spindle pulling force. On the other hand, Kip3 and Ndc80 showed a smilar linear localization in presence of the microtubules (DMSO, type II, Figure 3D,E) as observed before (Figure 2A, B) suggesting they are at all the kinetochores linearly aligned between two SPBs regardless of the microtubule attachment. Surprisingly, we observed that Kip3 also mostly dislodged from the kinetochore like Cin8 and Kip1 upon removal of the microtubules (Benomyl, type III, Figure 3D). However, Ndc80 showed a single bright dot with slight dispersed localization in absence of the microtubules (Benomyl, type III, Figure 3E) suggesting it was largely maintained at the homolog kinetochore clusters that coalesced due to lack of pulling force. In all the cases type I category with single dot-like localalization of the proteins depicts the cells that failed to enter into meiosis. These results indicate that the motors are dislodged from the centromeres more than Ndc80 upon removal of microtubules which is also supported by our ChIP assays (Figure 3F). Mislocalized Cin8 pool perhaps gets degraded as its overall intensity dropped with respect to Ndc80 upon treatment with benomyl (Figure 3G). Taken together, from the above results we conclude that Cin8 and Kip1 are stably maintained in a microtubule-dependent manner at the fully matured kinetochores that are attached end-on to the microtubules. On the other hand, Kip3 and Ndc80 are there at all the kinetochores with (end-on and lateral) or without microtubule attachment (Figure S5).

### Cin8 and Kip3 are required for stable localization of outer kinetochore protein Ndc80 at the centromeres

The outer kinetochore proteins are shredded by Ipl1 kinase activity at the beginning of meiosis negating kinetochore-microtubule attachment at early stage. This is a key event to facilitate monopolar attachment conferring the meiotic-specific chromosome segregation (Meyer et al., 2015a; Meyer et al., 2013). However, the determinant involved in the subsequent reassembly of the outer kinetochore proteins to form a mature kinetochore complex is unclear. Since we observed that the appearance of Cin8 and Kip1 at the kinetochores following prophase I coincided with the recruitment of Ndc80 at that site during metaphase I (Figure 1), we argued that the motors perhaps influence the reassembly of the outer kinetochore proteins. To examine this possibility, we investigated the localization of Ndc80-EGFP in the wild-type and the motor mutants in the metaphase I arrested cells. Compared to the wild-type and *kip1*Δ mutant, we observed a reduction in Ndc80 intensity in *cin8*Δ mutant and, to a lesser extent, in *kip3*Δ (Figure 4A and S3A).

**Figure 4:**
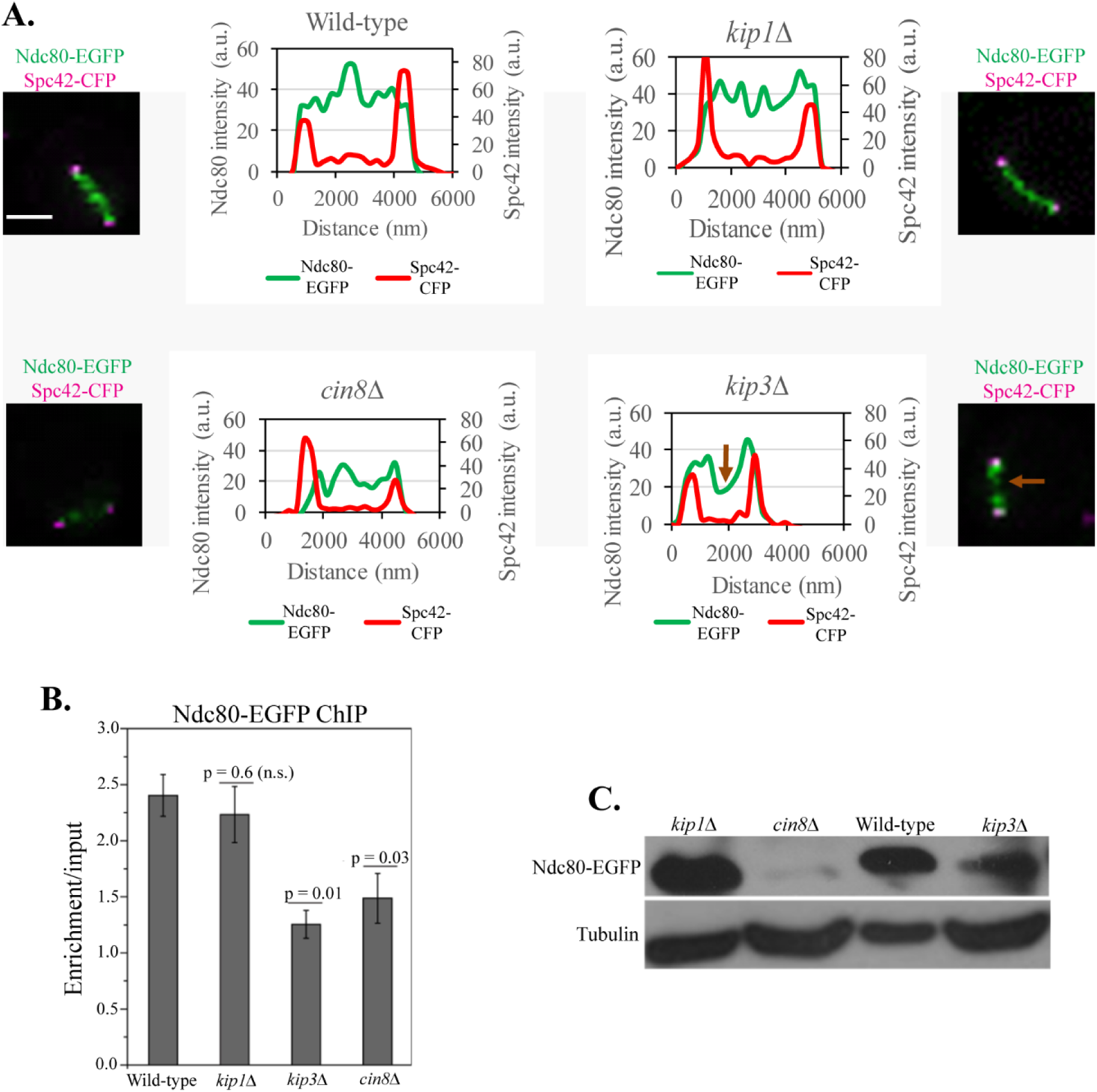
The absence of Cin8 and Kip3 reduces Ndc80 localization at the centromeres. (A) Representative images showing the localization of Ndc80-EGFP and the intensity plots of the same in the wild-type (SGY5313), *cin8*Δ (SGY5379), *kip1*Δ (SGY5423), and *kip3*Δ (SGY5424) metaphase I arrested cells (after 8 h of meiotic induction). Spc42-CFP was used as a marker to judge the metaphase spindle and as an internal control for the intensity measurement. The area under the graph represents Ndc80 intensity value obtained after normalization by the background fluorescence intensity as described in the materials and methods (B) ChIP assay showing association of Ndc80 with *CENIII* in the metaphase I arrested (as in A) wild-type (SGY5247), *cin8*Δ (SGY5248), *kip1*Δ (SGY5193), and *kip3*Δ (SGY5196) cells harbouring Ndc80-EGFP. The enrichment values of Ndc80 at *CENIII* were plotted. The error bar represents the SEM obtained from at least two independent replicates. p value > 0.05 were counted as not significant (n.s.). Bar, 2 µm. (C) Strains used in (A) were analysed for the degradation of endogenous Ndc80-EGFP in metaphase I arrested stage.

Notably, in contrast to *cin8*Δ where Ndc80 distribution was affected across the entire length of the spindle axis, in *kip3*Δ the reduction was prominent only at the mid-region between the two SPBs (Figure 4A and S3A). Consistent with this, when the total intensities of Ndc80-EGFP obtained from the wild-type and the mutant cells were plotted and compared, we observed a significant reduction in the signal intensity in *cin8* Δ and *kip3*Δ as compared to the wild-type and *kip1*Δ mutant (Figure S3B). The reduced signal of Ndc80 in *cin8*Δ or *kip3*Δ was further corroborated by quantitative ChIP assay from the metaphase I arrested cells, where we observed a reduced Ndc80 enrichment at the centromeres in *cin8* Δ and *kip3* Δ but not in *kip1*Δ (Figure 4B). Since kinetochore unbound Ndc80 eventually gets degraded over time (De Wulf *et al*., 2003), we could hardly detect any Ndc80 in *cin8* Δ, and a considerable reduction in its level was observed in *kip3* Δ cells that were arrested at metaphase I for prolonged 8 h duration (Figure 4C). As expected, the Ndc80 level remained similar in the wild-type and *kip1* Δ cells. The gross reduction of the Ndc80 signal in *cin8*Δ or *kip3*Δ cells is not due to lowering its expression level in these mutants under normal cycling condition (Figure S3C-D). These results suggest that Cin8 and, to some extent, Kip3 are required for stable localization of Ndc80 at the centromeres and thus contribute to reassembly of mature kinetochores at then beginning of meiosis.

In mitosis, earlier reports have demonstrated that Cin8 and Kip1 but not Kip3 localization at the centromeres is Ndc80 dependent (Tytell and Sorger, 2006). To examine if, similar to meiosis, the localization of Ndc80 to the centromere is dependent on the motor proteins during mitosis, we compared the intensity of Ndc80-EGFP between the wild-type and the motor mutants in metaphase cells as judged by the distance between the two SPBs. However, we failed to observe any significant difference (Figure S4A-C) which was consistent with the ChIP results (Figure S4D). Notably, in *cin8*Δ and *kip3*Δ cells, Ndc80-EGFP appeared linearly dispersed in contrast to bi-lobed appearance in the wild-type and *kip1*Δ cells, as reported earlier (Tytell and Sorger, 2006; Gardner *et al*., 2008; Wargacki *et al*., 2010). Therefore, we conclude that the role of motor proteins in properly depositing Ndc80 at the kinetochore leading to its maturation is meiosis-specific.

### The absence of Cin8 also affects centromeric maintenance of the inner kinetochore protein Ndc10

We and others have earlier reported that the inner and outer kinetochore proteins are interdependent for centromeric localization (Thakur and Sanyal, 2012; Mittal *et al*., 2019) in contrast to the notional hierarchical organization of the kinetochore at the centromeres. Therefore, it is plausible that the improper localization of outer kinetochore protein (Ndc80) at the centromeres in the motor mutant may hinder the maintenance of inner kinetochore proteins at the centromeres. To test this possibility, we measured the intensity of the inner kinetochore protein Ndc10 fused to EGFP in metaphase I arrested cells and observed a significant reduction of the intensity in the *cin8*Δ cells (Figure 5A). The reduction in the Ndc10-EGFP fluorescence signal in the *cin8*Δ cells prompted us to investigate its occupancy at the centromeres in the wild type and the motor mutants by ChIP assays. We observed a significant drop in Ndc10-EGFP level at *CENIII* only in *cin8*Δ but not in the other mutants (Figure 5B). However, there was no change in Ndc10-EGFP expression between the wild type and *cin8*Δ mutant (Figure 5C). These results indicate that the loss of Cin8 affects the assembly of the outer kinetochore, and the absence of outer proteins in right stochiomemtry perhaps affects the kinetochore ensemble in the long run causing dislodging of the inner kinetochore proteins.

**Figure 5:**
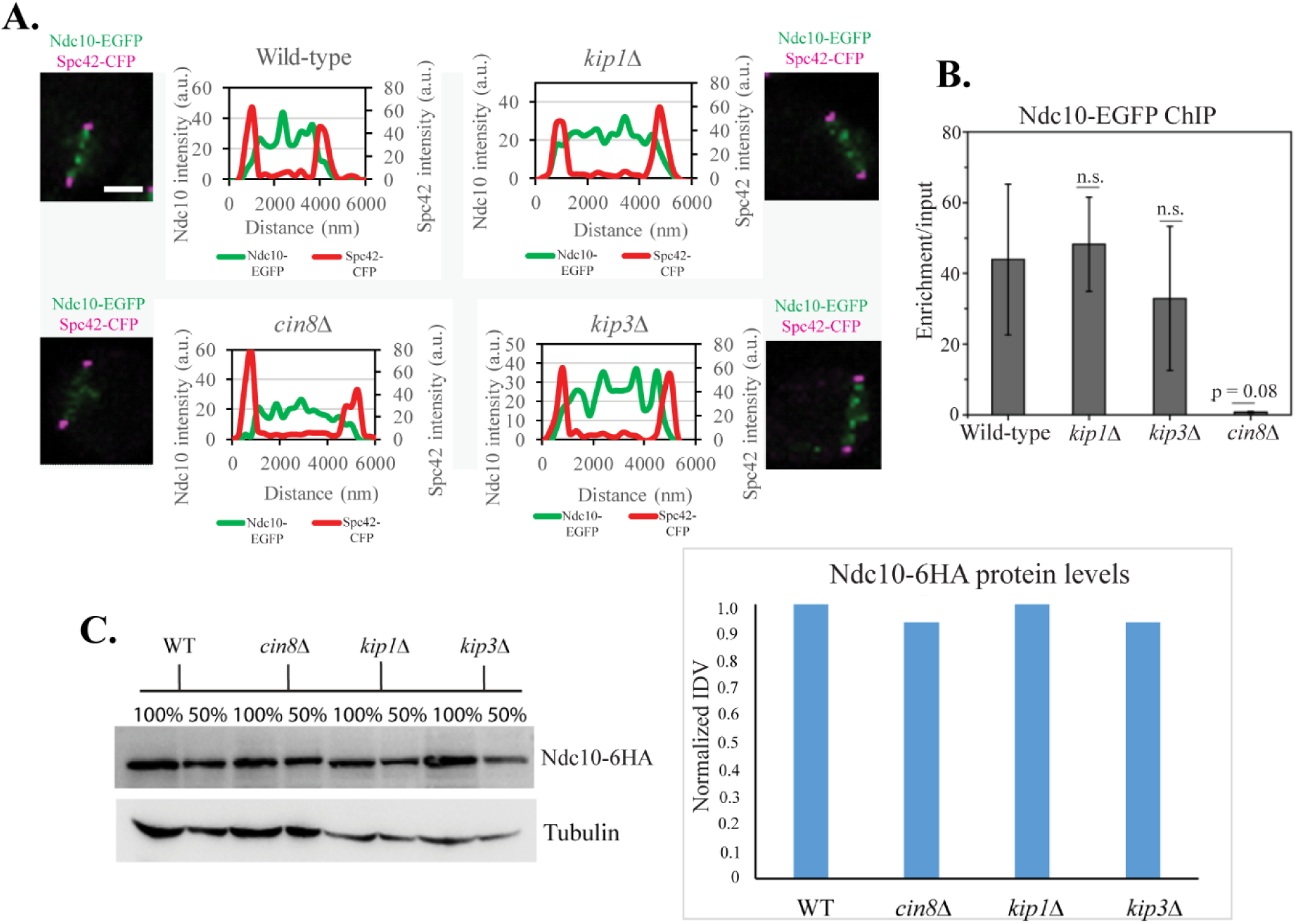
The absence of Cin8 perturbs centromeric maintenance of the inner kinetochore protein Ndc10. (A) Representative images showing the localization of Ndc10-EGFP and the intensity plots of the same in the wild-type (SGY5520), *cin8*Δ (SGY5519), *kip1*Δ (SGY5490), and *kip3*Δ (SGY5521) metaphase I arrested cells (after 8 h of meiotic induction). Spc42-CFP was used as a marker to judge the metaphase spindle and as an internal control for the intensity measurement. The area under the graph represents Ndc10 intensity value obtained after normalization by the background fluorescence intensity values as described in materials and methods. (B) ChIP assay showing association of Ndc10 with *CENIII* in the metaphase I arrested (as in A) wild-type (SGY5398), *cin8*Δ (SGY5383), *kip1*Δ (SGY5447), and *kip3*Δ (SGY5455) cells harbouring Ndc10-EGFP after 8 h of meiotic induction. The enrichment values of Ndc10 at *CENIII* were plotted. The error bar represents the SEM obtained from at least two independent replicates. p value > 0.05 were counted as not significant (n.s.). Bar, 2 µm. (C) Left, the expression levels of Ndc10-in the wild-type (SGY344), *cin8*Δ (SGY13021), *kip1*Δ (SGY13022), and *kip3*Δ (SGY13023) metaphase I arrested cells. Right, the intensity of the Ndc10-6HA bands corresponding to 50% dilution were compared and normalized with the tubulin bands.

## Discussion

Accurate chromosome segregation depends on stable KT-MT attachment, the establishment of which differs in a temporal fashion between the early stages of mitosis and meiosis in budding yeast. In mitosis while the cells are in S phase, the kinetochores are detached briefly during centromere replication and reattached immediately to the microctubules due to their rapid disassembly-reassembly (Tanaka et al 2005). In contrast, cells following entry into meioisis shed the outer kinetcohores but do not reassemble them immdeiately after centromere replication but do so much later during prophase I. The mechanisms engaged to resist reassembly of the kinetochores till prophase I are known (Chen et al., 2017; Chia et al., 2017) but the determinants that promote reassembly of the kinetcohores and reestablishment of stable KT-MT attachment are not fully known. Kinesin motors have been studied extensively in mitosis describing their myriad functions related to KT-MT attachment and microtubule spindle assembly-disassembly (Grissom et al., 2009; Gupta Jr et al., 2006; Mayr et al., 2007, Tanaka et al., 2007Gardner et al.,2008, Tytell & Sorger 2006, Shapira et al., 2017, Suzuki et al 2018). In those studies it was shown that outer kinetochore proteins and kinetochore microtubules recruit the kinesin motors at the centromeres to control KT-MT attachment. In this report we reveal that the kinesin motors are also involved in stabilization of the kinetochore ensemble and thus contributes to reestablishment of KT-MT attachment during early meiosis.

### The congression of the kinetochores towards spindle poles during metaphase I is less synchronous in meiosis than in mitosis

The observation that unlike mitosis, the characteristic ‘bi-lobed’ organization of all the kinetochores is absent during metaphase I in meiosis (Figure 2A), is somewhat surprising. This is because such an organization happens in mitotic metaphase due to bi-orientation of the sister kinetochores and is also expected to occur during metaphase I in meiosis as the homologs become bi-oriented. However, the presence of linearly disperesed kinetochores in between two SPBs (Figure 2A) with a similar localization of Kip3 (Figure 2B) incidates that i) the congression of the kinetochores towards the spindle poles is not synchronous and ii) Kip3 decorates all the kinetochores. This is in contrast with mitosis where the sister-kinetochore pairs congress almost together towards opposite spindle poles forming bi-lobed organization. We believe this happens as in mitosis all the kinetochores remain clustered and attached to the microtubules and following detachment for a very brief period during replication, they reestablish stable attachment with the microtubules almost together because they remain close to each other. However, in meiosis the kinetochroes without outer protein (Ndc80) till prophase I (Figure 1A) cannot connect to the microtubules and become declustered (Figure S1). Following prophase I as the kinetochores are reassembled, the individual dispersed kinetochore attaches to the microtubules in a stochastic manner. During this course of time an unattached kinetochore first establishes a unstable lateral attachment followed by a stable end-on attachment to the microtubules. We believe that among the linearly dispersed kinetochores, the ones that remain close or distal to the poles are attached end-on or lateral to the microtubules, respectively. This is supported by the localization patterns of Dam1 with respect to Ndc80 (Figure 3H). Kip3 localization data (Figure 2B) suggests that Kip3 can occupy both laterally and end-on attached kinetochores (Figure S5). On the other hand, the localization of the Cin8 and Kip1 only next to the SPBs in metaphase I (Figure 2B) argues that they occupy a subset of the kinetochores that are stably end-on attached and remain close to the SPBs (Figure S5). This is corroborated by the observations that i) Kip3 was found enriched at the centromeres compared to Cin8 and Kip1 at prophase I when kinetochore-microtubule attachment (Ndc80) is absent (Figure 1F) and ii) at the population of centromeres with reassembled kinetochores in metaphase I, the enrichment of Kip3 is more than Cin8 or Kip1 (Figure 3A). This is supported by the studies in mitosis where centromeric localization of Kip3 is found microtuble independent (Tytell & Sorger 2006) and Cin8/Kip1 localization depends on microtubules as well as on the presence of outer kinetochores (Tytell & Sorger 2006, Suzuki et al 2018). Although the metaphase I localization pattern suggests that Cin8/Kip1 and Kip3 perhaps differ in residing at the centromeres depending on the type of KT-MT attachment, we observed that during prolonged metaphase I arrest, the removal of microtubules destabilize centromeric localization of all the motors but not Ndc80 (Figure 3B-E, F, G) which is consistent with previous report from mitotic cells (Suzuki et al 2018). This suggests that in absence of microtubule attachment the outer kinetochore can no longer stabilize the Cin8/Kip1 and Kip3 kinesin motors. This may be in contrast with the other minus end directed kinesin motor, Kar3 that can occupy microtubule unattached kinetochore (Tytell & Sorger 2006, Tanaka et al., 2005).

### Kinesin-5 motor Cin8 contributes to reassemble stable kinetochores during early meiosis

Our cell biological and biochemical data show that during early meiosis Cin8 appears at the centromeres along with the outer protein, Ndc80 (Figure 1A, 1F, 2B, 3A). This is consistent with the earlier report that Cin8’s centromeric localization depends on Ndc80 complex (Suzuki et al., 2018). Interestingly, in this report we reveal that Cin8 in turn also promotes the stabilization of the kinetochores (marked by outer Ndc80 and inner Ndc10 proteins) as they reassemble following prophase I (Figure 4, Figure 5). Although we noticed a similar decrease in Ndc80 at the centromeres in absence of Kip3 as well (Figure 4B) but that decrease happens at the centromeres that are at the middle of the spindle axis (shown by arrow in Figure 4A, *kip3*Δ graph) but not at the SPB proximal and end-on attached kinetochores as observed in *cin8*Δ cells (Figure 4A, *cin8*Δ graph). The reduction of Ndc10 level at the centromeres in absence of Cin8 (Figure 5A) perhaps happens indirectly due to perturbation in centromeric maintenance of Ndc80 as we and other groups have shown earlier that in yeast inter-kinetochore interactions are important to maintain kinetochore integrity (Richmond et al., 2013, Mittal et al., 2019, Thakur & Sanyal 2012).

Due to the difference in chromosome segregation patterns between mitosis and meiosis, several kinetochore proteins have altered essentiality (Mehta et al., 2014, Agarwal et al., 2015). Likewise, earlier we reported absence of Cin8 and Kip3 severely compromises genome stability specifically in meiosis (Mittal et al., 2019). Here, we add to that list by showing a meiosis-specific kinetochore stabilizing function of the kinesin motors. In various higher eukaryotes particularly with open nuclear divisions, including humans, the kinesin and dynein motors are shown to be involved in gradual establishment of stable KT-MT attachment through conversion of lateral to end-on attachments (Vaisberg et al., 1993, Shrestha et al., 2013). In this work, we could unravel the similar functions of kinesins 5 in yeast with closed nuclear division as in meiosis the KT-MT attachment dynamics is similar to the open nuclear system. Taken together, we provide the first evidence of the importance of the motor proteins in stabilization of the kinetochore ensemble during meiosis in budding yeast. Many studies in myriad systems have shown that kinetohcores recruit motor proteins at the centromeres which in turn modulate the dynamics of KT-MT attachment. But to our knowledge, whether motors in turn can stabilize the kinetochore has not been shown in any system regardeless of the type of cell cycle. It is likely that Cin8’s role in stabilization of the kinetochore is actually through stabilization of plus-ends of the microtubule within the kinetochores via end-on attachment. But a microtubule independent kinetochore stabilization function through an interaction between Ndc80 complex and Cin8 cannot be ruled out. Although Cin8 has been found not to interact with Ndc80 in vitro (Suzuki et al., 2018), but its in vivo interaction with Nuf2 (Newman et al., 2000) argues for a stability promoting interaction with Ndc80 complex. Future studies in metazoans would tell whether such meiosis-specific kinetochore stabilization role of the kinesin motors is conserved across eukaryotes.

## Abbreviations

KT-MT: kinetochore-microtubule
SPB: spindle pole body
SAC: Spindle assembly checkpoint

## Acknowledgments

We are thankful to the central instrumental facility of IIT Bombay for the laser scanning confocal microscope. The laboratory of the SKG is supported by Department of Biotechnology, Govt. of India (BT/PR20932/BRB/10/1539/2016) and Science and Engineering Research Board, Govt. of India (CRG/2020/000444). SS is supported by a DBT fellowship (DBT/2018/IIT-B/1062), PM is supported by UGC fellowship (17-06/2012(i) EU-V) DK is supported by a DBT fellowship (DBT/2018/IIT-B/1059) and AM is supported by IITB-Monash Research Academy, India (IMURA0907).

---

**Figure S1:**
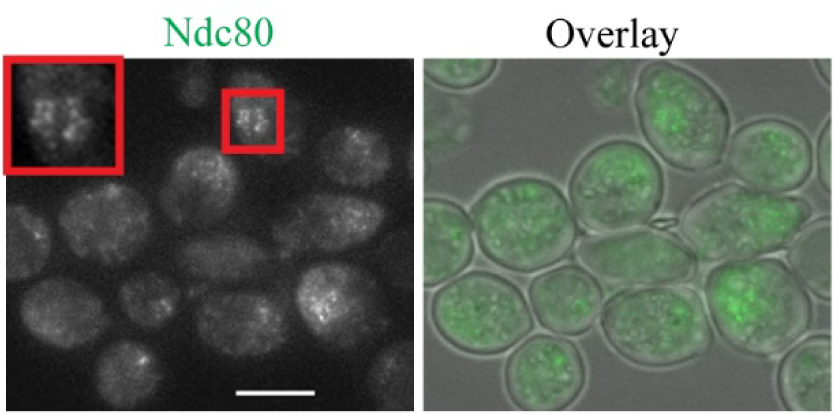
Declustered localization of Ndc10-EGFP in prophase I. The strain harboring Ndc10-EGFP (SGY5351) was induced for meiosis in sporulation medium (SPM) for 6 h and was arrested at prophase I by Ndt80 depletion. The arrested live cells were imaged through fluorescence microscopy. Bar, 5 µm.

**Figure S2:**
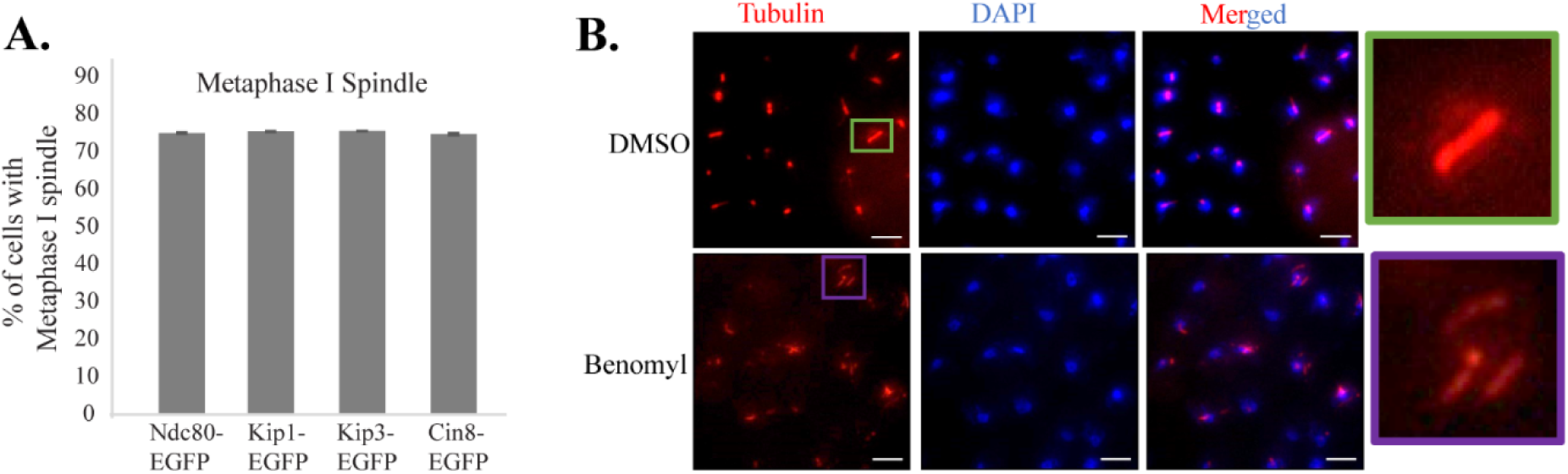
Benomyl treatment of cells arrested at metaphase I. (A) Ndc80-EGFP (SGY5247), Kip1-EGFP (SGY5117), Kip3-EGFP (SGY5348), and Cin8-EGFP (SGY5119) cells were arrested at metaphase I by Cdc20 depletion using p*CLB2* promoter. (n > 100 cells for each strain). Tubulin immunostaining was done using anti-tubulin antibodies to count the cells with metaphase I spindle. The error bar represents SEM obtained from two replicates. (B) Tubulin immunostaining to confirm the microtubule depolymerization following DMSO or benomyl treatment of the Cdc20-arrested cells used in A. Bar, 5 µm

**Figure S3:**
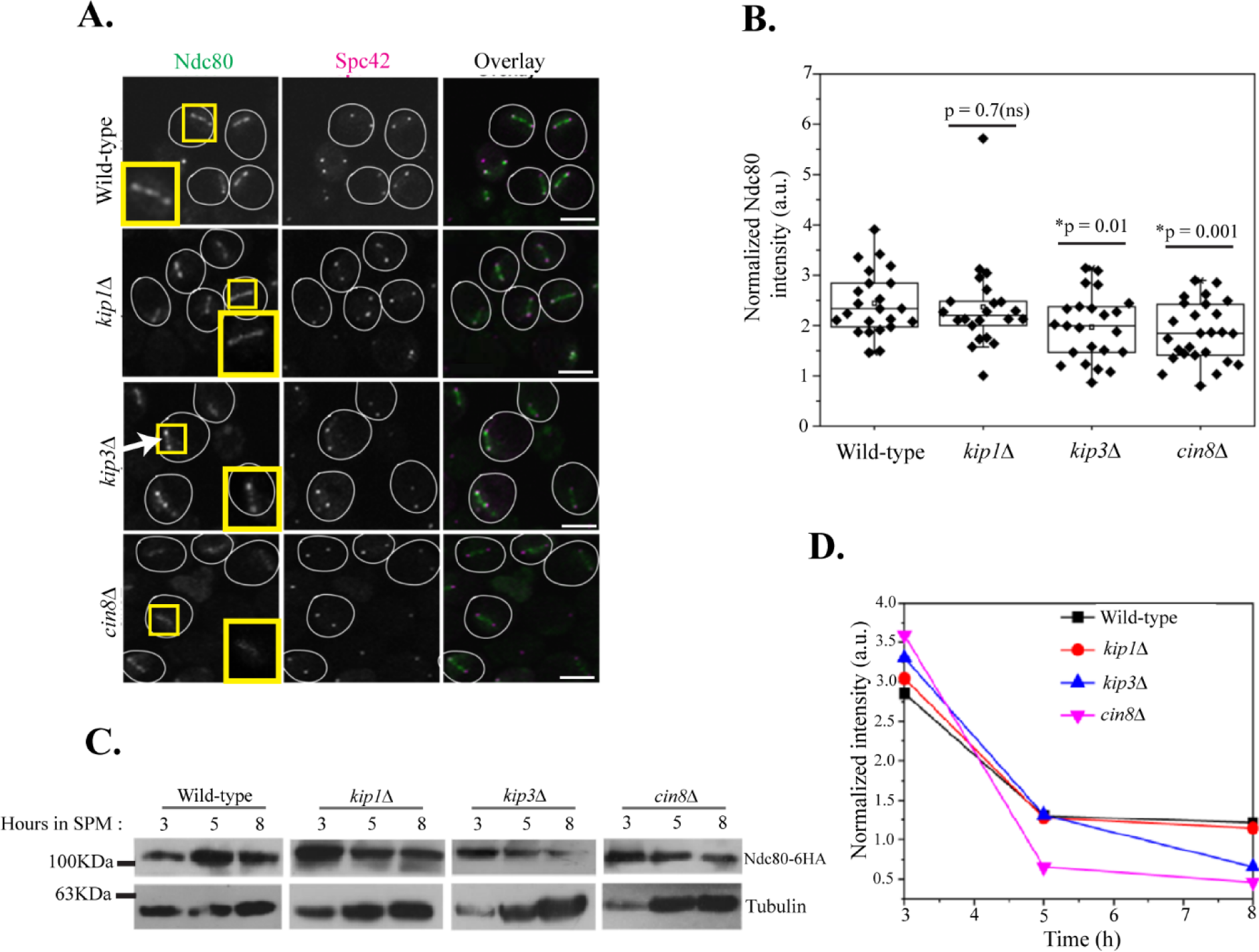
Localization and expression level of Ndc80 in the wild-type and motor mutants. (A) Expanded field views of the strains used in figure 4A show several cells with reduced localization of Ndc80 in *cin8*Δ and *kip3*Δ than in the wild-type and *kip1*Δ. Arrow indicates the drop in the Ndc80 localization in the middle region of the two SPBs observed in *kip3*Δ cells. Images were acquired after 8 h of incubation in the sporulation medium. (B) Quantification of Ndc80-EGFP fluorescence intensity for the strains used in figure 4A after normalization with the background and Spc42-CFP intensity (internal control) value in the respective cell. (C) Ndc80 expression is not altered in the motor mutants. Immunoblotting was used to detect Ndc80-6HA from the wild-type (SGY 372), *cin8*Δ (SGY5469), *kip1*Δ (SGY5467), and *kip3*Δ (SGY5468) cells. For protein extraction, the cells were harvested following induction of meiosis for the indicated h. The band obtained for α-tubulin served as a loading control. (D) Quantification of Ndc80-6HA bands at different time points from the blot in (C) after normalization using the background and α-tubulin band intensities. Bar, 5 µm.

**Figure S4:**
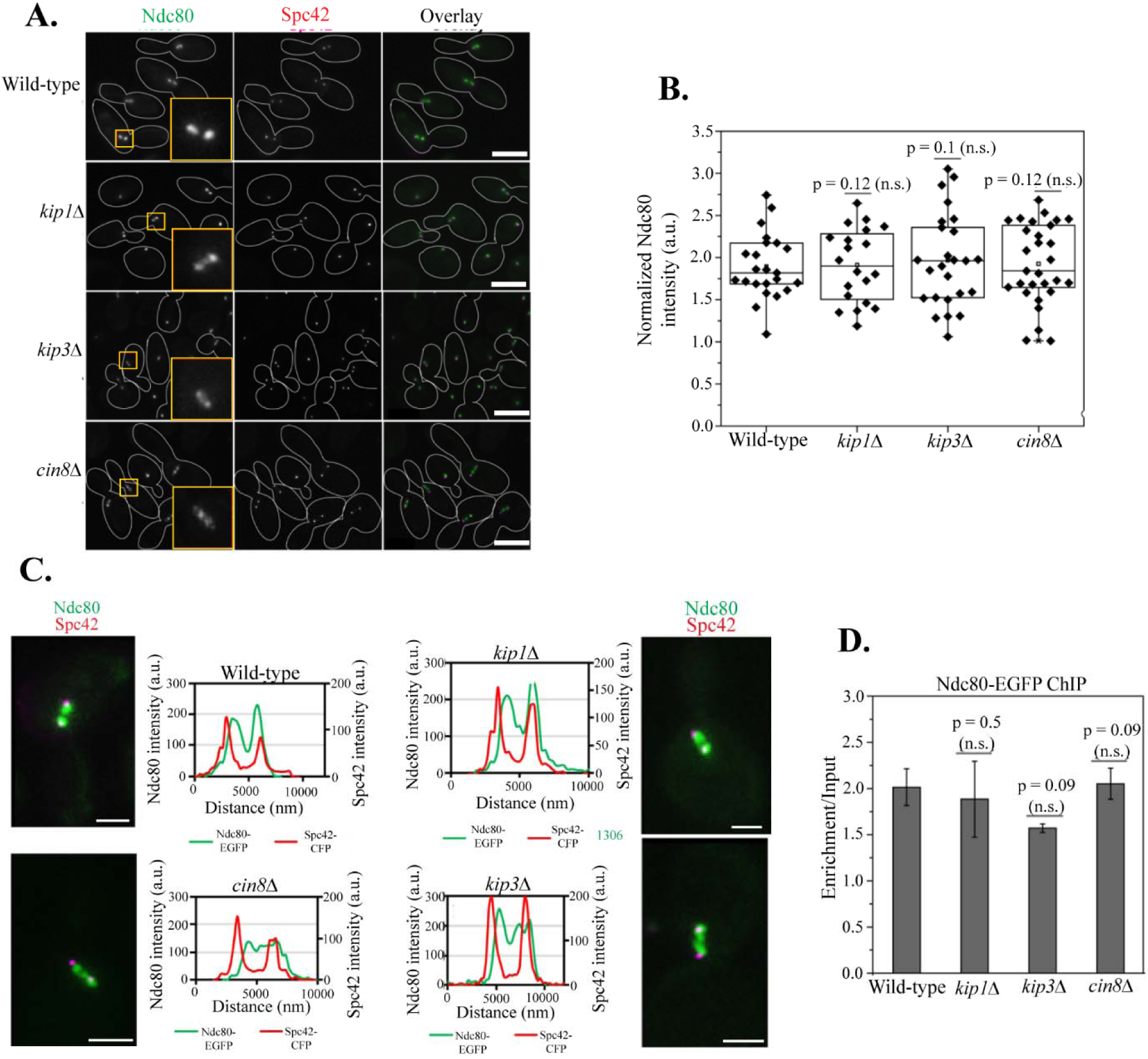
Localization and level of Ndc80 in the wild-type and motor mutants in mitotic metaphase cells. (A) Expanded field views of the cells showing Ndc80-EGFP and Spc42-CFP in wild-type (SGY5313), *cin8*Δ (SGY5379), *kip1*Δ (SGY5423), and *kip3*Δ (SGY5424) arrested in mitotic metaphase using auxin mediated Cdc20 depletion. Bar, 5 µm. (B) Quantification of Ndc80-EGFP fluorescence intensity for the strains used in (A) after normalization with the background and Spc42-CFP intensity (internal control) value in the respective cell. (C) Representative images showing the localization of Ndc80-EGFP and Spc42-CFP and their intensity plots from the strains used in (A). The area under the graph represents Ndc80 intensity value obtained after normalization by the total Spc42 intensity values as described in the materials and methods. Bar, 2 µm. (D) ChIP assay showing association of Ndc80 with *CENIII* locus in the metaphase arrested of the strains used in (A). The enrichment values of Ndc80 with *CENIII* were plotted. The error bar represents the SEM obtained from at least two independent replicates. p value > 0.05 were counted as not significant (n.s.). Bar, 2 μm. (C) Strains used in (A) were analysed for the degradation of endogenous Ndc80-EGFP in metaphase I arrested stage.

**Figure S5:**
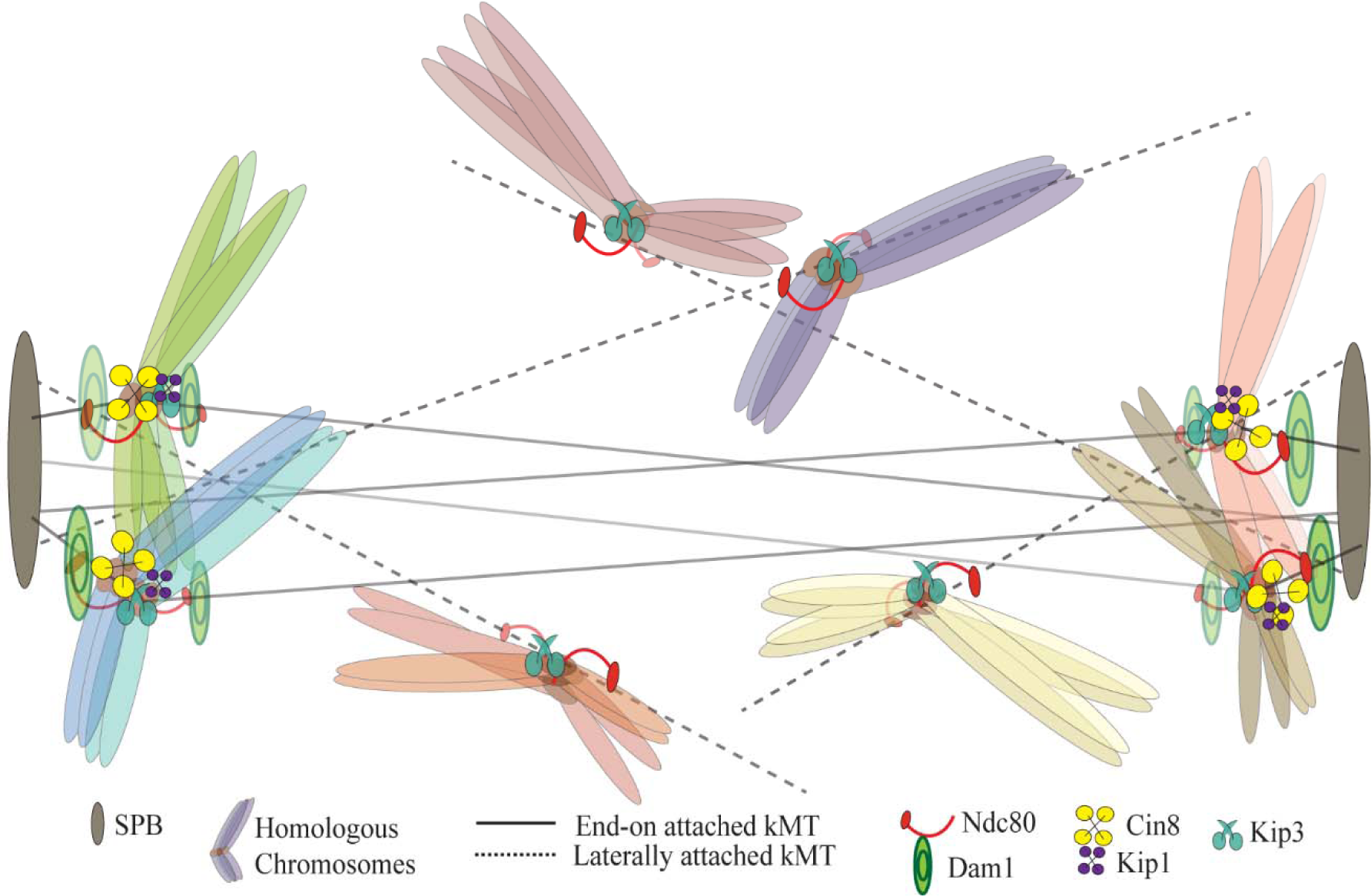
A working model of the kinetochore stabilization. A schematic representation of congressing chromosomes during metaphase I depicting the localization of Cin8, Kip1, and Dam1 at the kinetochores that are attached end-on (solid lines) to the microtubule and remain close to the SPBs; the localization of Kip3 and Ndc80 at the kinetochores that are either unattached (not shown), laterally (dashed lines) or end-on attached to the microtubules.

## Notes

### Competing Interest Statement

The authors have declared no competing interest.

