## Supplemental Tables for "Evidence of kinesin motors involved in stable kinetochore assembly during early meiosis"

List of yeast strains used in this study:

All the strains were derived from parent haploid strains of SK1 background with the following genotypes.

MATa, ho::Lys2, lys2, ura3, leu2::hisG, his3::hisG, trp1::hisG

MATα, ho::Lys2, lys2, ura3, leu2::hisG, his3::hisG, trp1::hisG

**Table 1: List of Strains**

| **Yeast strain** | **Genotype** |
| --- | --- |
| SGY344 | *MATa/α*, *cdc20::pCLB2-CDC20::KANMX6/ cdc20::pCLB2 CDC20::KANMX6, NDC10-6HA::HIS3/NDC10-6HA::HIS3* |
| SGY372 | *MATa/α*, *cdc20::pCLB2-CDC20::KANMX6/ cdc20::pCLB2-CDC20::KANMX6, NDC80-6HA::HIS3/NDC80-6HA::HIS3* |
| SGY4203 | *MATa/α*, *cdc20::pCLB2-CDC20::KANMX6/ cdc20::pCLB2-CDC20::KANMX6, CIN8-6HA::HIS3/CIN8-6HA::HIS3* |
| SGY4206 | *MATa/α*, *cdc20::pCLB2-CDC20::KANMX6/ cdc20::pCLB2-CDC20::KANMX6, NDC80-6HA::HIS3/NDC80-6HA::HIS3* |
| SGY5051 | *MATa/α*, *KIP1-EGFP::TRP1/KIP1-EGFP::TRP1, NDC80-CFP::HIS3/NDC80-CFP::HIS3, SPC42-::KANMX6/SPC42-mCherry::KANMX6* |
| SGY5070 | *MATa/α*, *CIN8-EGFP::TRP1/CIN8-EGFP::TRP1, NDC80-CFP::HIS3/NDC80-CFP::HIS3, SPC42-mCherry::KANMX6/SPC42-mCherry::KANMX6* |
| SGY5109 | *MATa/α*, *ndt80::pGAL-NDT80::TRP1/ndt80::pGAL-NDT80::TRP1, CIN8-EGFP::TRP1/CIN8-EGFP::TRP1* |
| SGY5117 | *MATa/α*, *cdc20::pCLB2-CDC20::KANMX6/ cdc20::pCLB2-CDC20::KANMX6, KIP1-EGFP::TRP1/KIP1-EGFP::TRP1* |
| SGY5119 | *MATa/α*, *cdc20::pCLB2-CDC20::KANMX6/ cdc20::pCLB2-CDC20::KANMX6, CIN8-EGFP::TRP1/CIN8-EGFP::TRP1* |
| SGY5130 | *MATa/α*, *NDC80-EGFP::TRP1/NDC80-EGFP::TRP1, SPC42- mCherry::KANMX6/SPC42- mCherry::KANMX6* |
| SGY5151 | *MATa/α*, *ndt80::pGAL-NDT80::TRP1/ndt80::pGAL-NDT80::TRP1*, *CIN8-EGFP::TRP1/CIN8-EGFP::TRP1, SPC42-mCherry::KANMX6/ SPC42- mCherry::KANMX6, NDC80-CFP::HIS3/NDC80-CFP::HIS3* |
| SGY5165 | *MATa/α*, *KIP3-EGFP::TRP1/KIP3-EGFP::TRP1, NDC80-CFP::HIS3/NDC80-CFP::HIS3, SPC42- mCherry::KANMX6/SPC42- mCherry::KANMX6* |
| SGY5193 | *MATa/α*, *cdc20::pCLB2-CDC20::KANMX6/ cdc20::pCLB2-CDC20::KANMX6, NDC80-EGFP::TRP1/NDC80-EGFP::TRP1, kip1∆::URA3/ kip1∆::URA3* |
| SGY5196 | *MATa/α*, *cdc20::pCLB2-CDC20::KANMX6/ cdc20::pCLB2-CDC20::KANMX6, NDC80-EGFP::TRP1/NDC80-EGFP::TRP1, kip3∆::URA3/ kip3∆::URA3* |
| SGY5198 | *MATa/α, NDC10-EGFP::TRP1/ NDC10-EGFP::TRP1, NDC80-CFP::HIS3/NDC80-CFP::HIS3, SPC42- mCherry::KANMX6/SPC42- mCherry::KANMX6* |
| SGY5220 | *MATa/α*, *ndt80::pGAL-NDT80::TRP1/ndt80::pGAL-NDT80::TRP1, KIP1-EGFP::TRP1/KIP1-EGFP::TRP1* |
| SGY5247 | *MATa/α*, *cdc20::pCLB2-CDC20::KANMX6/ cdc20::pCLB2-CDC20::KANMX6, NDC80-EGFP::TRP1/NDC80-EGFP::TRP1* |
| SGY5248 | *MATa/α*, *cdc20::pCLB2-CDC20::KANMX6/ cdc20::pCLB2-CDC20::KANMX6, NDC80-EGFP::TRP1/NDC80-EGFP::TRP1, cin8∆::URA3/ cin8∆::URA3* |
| SGY5256 | *MATa/α*, *ndt80::pGAL-NDT80::TRP1/ndt80::pGAL-NDT80::TRP1, KIP3-EGFP::TRP1/KIP3-EGFP::TRP1* |
| SGY5266 | *MATa/α*, *ndt80::pGAL-NDT80::TRP1/ndt80::pGAL-NDT80::TRP1, KIP1-EGFP::TRP1/KIP1-EGFP::TRP1, SPC42- mCherry::KANMX6/SPC42- mCherry::KANMX6, NDC80-CFP::HIS3/NDC80-CFP::HIS3* |
| SGY5272 | *MATa/α*, *ndt80::pGAL-NDT80::TRP1/ndt80::pGAL-NDT80::TRP1*, *KIP3-EGFP::TRP1/KIP3-EGFP::TRP1, SPC42-mCherry::KANMX6/ SPC42- mCherry::KANMX6, NDC80-CFP::HIS3/NDC80-CFP::HIS3* |
| SGY5313 | *MATa/α*, *cdc20::pCLB2-CDC20::KANMX6/ cdc20::pCLB2-CDC20::KANMX6, NDC80-EGFP::TRP1/NDC80-EGFP::TRP1, SPC42-CFP::HIS3/SPC42-CFP::HIS3* |
| SGY5348 | *MATa/α*, *cdc20::pCLB2-CDC20::KANMX6/ cdc20::pCLB2-CDC20::KANMX6, KIP3-EGFP::TRP1/KIP3-EGFP::TRP1* |
| SGY5351 | *MATa/α*, *ndt80::pGAL-NDT80::TRP1/ndt80::pGAL-NDT80::TRP1, NDC10-EGFP::HIS3/NDC10-EGFP::HIS3* |
| SGY5379 | *MATa/α*, *cdc20::pCLB2-CDC20::KANMX6/ cdc20::pCLB2-CDC20::KANMX6, NDC80-EGFP::TRP1/NDC80-EGFP::TRP1, SPC42-CFP::HIS3/SPC42-CFP::HIS3, cin8∆::LEU2/ cin8∆::LEU2* |
| SGY5383 | *MATa/α*, *cdc20::pCLB2-CDC20::KANMX6/ cdc20::pCLB2-CDC20::KANMX6, NDC10-EGFP::TRP1/NDC10-EGFP::TRP1, cin8∆::LEU2/ cin8∆::LEU2* |
| SGY5398 | *MATa/α*, *cdc20::pCLB2-CDC20::KANMX6/ cdc20::pCLB2-CDC20::KANMX6, NDC10-EGFP::TRP1/NDC10-EGFP::TRP1* |
| SGY5423 | *MATa/α*, *cdc20::pCLB2-CDC20::KANMX6/ cdc20::pCLB2-CDC20::KANMX6, NDC80-EGFP::TRP1/NDC80-EGFP::TRP1, SPC42-CFP::HIS3/SPC42-CFP::HIS3, kip1∆::URA3/ kip1∆::URA3* |
| SGY5424 | *MATa/α*, *cdc20::pCLB2-CDC20::KANMX6/ cdc20::pCLB2-CDC20::KANMX6, NDC80-EGFP::TRP1/NDC80-EGFP::TRP1, SPC42-CFP::HIS3/SPC42-CFP::HIS3, kip3∆::URA3/ kip3∆::URA3* |
| SGY5447 | *MATa/α*, *cdc20::pCLB2-CDC20::KANMX6/ cdc20::pCLB2-CDC20::KANMX6, NDC10-EGFP::TRP1/NDC10-EGFP::TRP1, kip1∆::LEU2/ kip1∆::LEU2* |
| SGY5455 | *MATa/α*, *cdc20::pCLB2-CDC20::KANMX6/ cdc20::pCLB2-CDC20::KANMX6, NDC10-EGFP::TRP1/NDC10-EGFP::TRP1, kip3∆::LEU2/ kip3∆::LEU2* |
| SGY5467 | *MATa/α*, *cdc20::pCLB2-CDC20::KANMX6/ cdc20::pCLB2-CDC20::KANMX6, NDC80-6HA::HIS3/NDC80-6HA::HIS3, kip1∆::URA3/ kip1∆::URA3* |
| SGY5468 | *MATa/α*, *cdc20::pCLB2-CDC20::KANMX6/ cdc20::pCLB2-CDC20::KANMX6, NDC80-6HA::HIS3/NDC80-6HA::HIS3, kip3∆::URA3/ kip3∆::URA3* |
| SGY5469 | *MATa/α*, *cdc20::pCLB2-CDC20::KANMX6/ cdc20::pCLB2-CDC20::KANMX6, NDC80-6HA::HIS3/NDC80-6HA::HIS3, cin8∆::URA3/ cin8∆::URA3* |
| SGY5490 | *MATa/α*, *cdc20::pCLB2-CDC20::KANMX6/ cdc20::pCLB2-CDC20::KANMX6, NDC10-EGFP::TRP1/NDC10-EGFP::TRP1, SPC42-CFP::HIS3/SPC42-CFP::HIS3, kip1∆::LEU2/ kip1∆::LEU2* |
| SGY5519 | *MATa/α*, *cdc20::pCLB2-CDC20::KANMX6/ cdc20::pCLB2-CDC20::KANMX6, NDC10-EGFP::TRP1/NDC10-EGFP::TRP1, SPC42-CFP::HIS3/SPC42-CFP::HIS3, cin8∆::URA3/ cin8∆::URA3* |
| SGY5520 | *MATa/α*, *cdc20::pCLB2-CDC20::KANMX6/ cdc20::pCLB2-CDC20::KANMX6, NDC10-EGFP::TRP1/NDC10-EGFP::TRP1, SPC42-CFP::HIS3/SPC42-CFP::HIS3* |
| SGY5521 | *MATa/α*, *cdc20::pCLB2-CDC20::KANMX6/ cdc20::pCLB2-CDC20::KANMX6, NDC10-EGFP::TRP1/NDC10-EGFP::TRP1, SPC42-CFP::HIS3/SPC42-CFP::HIS3, kip3∆:: LEU2/ kip3∆:: LEU2* |
| SGY5718 | *MATa/α, DAM1-EGFP::TRP1/ DAM1-EGFP::TRP1, NDC80-CFP::HIS3/NDC80-CFP::HIS3, SPC42- mCherry::KANMX6/SPC42- mCherry::KANMX6* |
| SGY13021 | *MATa/α*, *cdc20::pCLB2-CDC20::KANMX6/ cdc20::pCLB2-CDC20::KANMX6, NDC10-6HA::HIS3/NDC10-6HA::HIS3, cin8∆::URA3/ cin8∆::URA3* |
| SGY13022 | *MATa/α*, *cdc20::pCLB2-CDC20::KANMX6/ cdc20::pCLB2-CDC20::KANMX6, NDC10-6HA::HIS3/NDC10-6HA::HIS3, kip1∆::HPHMX6/ kip1∆::HPHMX6* |
| SGY13023 | *MATa/α*, *cdc20::pCLB2-CDC20::KANMX6/ cdc20::pCLB2-CDC20::KANMX6, NDC10-6HA::HIS3/NDC10-6HA::HIS3, kip3∆::HPHMX6/ kip3∆::HPHMX6* |

**Table 2: List of ChIP Primers**

| **Primer Name** | **Description** | **Sequence** |
| --- | --- | --- |
| GM91 | Chr*IV* Arm (1081215 to 1081234) | GTCGATGGTTTCATTCAGAT |
| GM92 | Chr*IV* Arm (1081288 to 1081307) | CAATGGAGAGAGTGGATGTT |
| GM107 | *CENIII* F primer (V93) | GATCAGCGCCAAACAATATGG |
| GM108 | *CENIII* R primer (V94) | AACTTCCACCAGTAAACGTTT |
| KD7 | *TUB2* locus forward primer | CTTGTAGACAGCGTCATGG |
| KD8 | *TUB2* locus reverse primer | CAGATGTCATAAAGTGCTTCG |
